# Loss of pyrethroid resistance in newly established laboratory colonies of *Aedes aegypti*

**DOI:** 10.1101/760710

**Authors:** Farah Z. Vera-Maloof, Karla Saavedra-Rodriguez, Rosa P. Penilla-Navarro, Americo D. Rodriguez-Ramirez, Felipe Dzul, Pablo Manrique, William C. Black

## Abstract

**Background:** Resistance to pyrethroid insecticides in *Aedes aegypti* has become widespread after almost two decades of their frequent use to reduce arbovirus transmission. Despite this, use of pyrethroids continues because they are relatively inexpensive and because of their low human toxicity. Resistance management has been proposed as a means to retain the use of pyrethroids in natural populations. A key component of resistance management assumes that there is a negative fitness associated with resistance alleles so that when insecticides are removed, resistance alleles will decline in frequency. At least three studies in *Ae. aegypti* have demonstrated a decrease in pyrethroid resistance once the insecticide is removed.

**Methods/Principal Findings:** The present study aims to evaluate variation in the loss of pyrethroid resistance among newly established laboratory populations of *Ae. aegypti* from Mexico. Eight field collections were maintained for up to eight generations and we recorded changes in the frequencies of mutations at the V1,016I locus and at the F1,534C locus in the voltage gated sodium channel (*VGSC*) gene. I1,016 and C1,534 confer resistance. We also examined resistance ratios (RR) with type 1 and 2 pyrethroids.

**Conclusions/Significance:** We demonstrate that, in general, the frequency of the *Ae. aegypti* pyrethroid resistance alleles I1,016 and C1,534 decline when they are freed from pyrethroid pressure in the laboratory. However, the pattern of decline is strain dependent. In agreement with earlier studies, RR was positively correlated with I1,016 resistant allele frequencies and showed significant protection against permethrin, and deltamethrin whereas F1534C showed protection against permethrin but not against deltamethrin.

**Author Summary:** The author is interested in the evolution of genes that confer resistance to insecticides, especially when this evolution affects binding of insecticides to their target site. The Voltage Gated Sodium Channel gene represents an excellent opportunity to understand how mutations at the target site(s) affect the evolution of resistance in many different pest insect species including *Aedes aegypti*, the primary vector of Dengue Virus, Yellow Fever, Zika and Chikungunya arboviruses.

## Introduction

After almost two decades of frequent pyrethroid use for *Aedes aegypti* (L.) control there is now widespread resistance [1, 2]. Despite this, use of pyrethroids continues because they are relatively inexpensive and because they have low human toxicity. Resistance management has been proposed as a means of retaining the use of pyrethroids in natural populations [2]. A key component of resistance management assumes that there will be a negative fitness associated with resistance alleles so that when insecticides are removed, resistance alleles will decline in frequency.

Laboratory strains of *Ae. aegypti* have shown a decrease in resistance once pyrethroid is removed suggesting a fitness cost associated with resistance. To date three laboratory studies have evaluated the loss of pyrethroid resistance in *Ae. aegypti*. In Taiwan, a permethrin resistant laboratory strain was maintained for 47 generations under permethrin pressure. Following 15 generations without exposure there was a significant decrease in permethrin resistance ratio (RR) and resistance alleles in the Voltage Gated Sodium Channel (VGSC) gene [3]. In Brazil, after 15 generations, the frequency of I1,016 decreased from 0.75 to 0.20 [4]. A study in Mexico, showed a significant increase in the proportion of knocked-down mosquitoes after 10 generations following removal of pyrethroids [5] but without a decrease in the frequency of I,1016 and C1,534 mutations in the VGSC. All three studies were done in laboratory cages.

The present study aims to evaluate the loss of pyrethroid resistance from eight collections of *Ae. aegypti*, (six field collections from or near the city of Merida and two collections from Tapachula and Acapulco from southern Mexico). These collections were maintained without pyrethroid pressure for eight consecutive generations during which we recorded changes in the frequencies of the two VGSC mutations I1,016 and C1,534, and the analysis of resistance ratios (RR) with permethrin (pyrethroid type 1) and deltamethrin (pyrethroid type 2).

## Materials and Methods

### *Aedes aegypti* field populations

In 2014, larvae were collected from eight public sites in Mexico. These were reared to adults and identified as *Ae. aegypti*. These were then blood fed (citrated sheep blood – Colorado Serum Co., Denver Colorado) in an artificial membrane feeder and eggs were collected for shipment. The GPS coordinates and name abbreviations of the collection sites appear in Table 1. Eggs from Yucatan were collected from three sites in urban areas of Merida and three collections from villages near Merida. Two additional collections were made in Tapachula and Acapulco in the states of Chiapas and Guerrero, respectively.

**Table 1.** States and cities of collection sites, geographical coordinates and site’s abbreviations used in this study.

| State | City | Site | Latitude | Longitude | Abbreviation |
| --- | --- | --- | --- | --- | --- |
| **Guerrero** |  |  |  |  |  |
|  | Acapulco | Zapata | 16.9049222 | -99.8410944 | **Acp** |
| **Chiapas** |  |  |  |  |  |
|  | Tapachula | Col.5 de Febrero | 14.9204944 | -92.2593472 | **Tap** |
| **Yucatan** |  |  |  |  |  |
|  | Merida | Fco. Montejo | 21.0307194 | -89.6463639 | **Mer1** |
|  |  | Plan Ayala | 21.0135833 | -89.6222222 | **Mer2** |
|  |  | U.H. Morelos | 20.9420139 | -89.5981556 | **Mer3** |
|  | Conkal | Center | 21.0747917 | -89.5199056 | **Co** |
|  | Dzitya | Center | 21.0623278 | -89.6746694 | **Dz** |
|  | Acanceh | Center | 20.8126083 | -89.4505611 | **Ac** |

### Establishment and maintenance of field populations

F_1_ eggs were sent to Colorado State University. Egg papers were placed into a water container with 2 L of tap water to promote development and hatching. Larvae were fed 2 mL of 10% (w/v) liver powder every other day. We transferred pupae to plastic cages for adult emergence. Larvae and mosquitoes were maintained in an incubator at 27-28°C, 70-80% humidity, and a photoperiod of 12h light:12h dark. Adults were fed with raisins and allowed access to tap water. Females were offered citrated sheep blood on artificial membrane feeders, every four days, to obtain eggs. Females laid their eggs on moistened filter paper. The eggs were allowed to mature for 48 hr before they were partially dried at room temperature and stored in sealed plastic bags. Each collection was split at the F_1_ larval stage into three groups of 50 to act as three biological replicates. In each subsequent generation ~25 adult ♀ and 25 ♂ were used to maintain each of the three replicates.

### Genotyping V1,016I and F1,534C

DNA was extracted, at each generation (F_1_-F_8_), from individual mosquitoes by the salt extraction method [6] and suspended in 180 µL of TE buffer. To identify allelic variation we used the allele-specific polymerase chain reaction (asPCR), followed by generation of a melting curve (CFX-96 BioRad), to identify genotypes [7–9]. In each of the eight generations, we analyzed three replicates of 50 adult mosquitoes (~25♀ and 25♂) for each of the eight collection sites. Sample sizes were kept intentionally large to minimize founder’s effects and genetic drift.

### Allele frequencies and linkage disequilibrium analysis

We estimated allele frequencies in each of the eight generations from the genotypic frequencies ((resistant allele frequency = ((2*resistant homozygote) + heterozygote)) / (2*sample size)). Allele frequencies were compared among replicates using a 2×3 contingency χ^2^ test. WINBugs2.0 [10] with 10^6^ iterations was used to calculate the 95% high-density intervals (HDI 95%) around allele and genotype frequencies. We used ggplot2 in R-3.5.1 to graph the data. In addition, we used LINKDIS [11] and χ^2^ tests to calculate the pairwise linkage disequilibrium coefficients (R_ij_) between alleles at loci 1,016 and 1,534 [12] where:

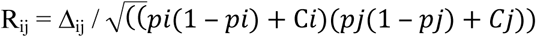

where 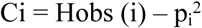 and Hobs(i) is the observed frequency of i homozygotes and

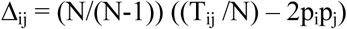

N is the sample size and p_i_ and p_j_ are the frequencies of alleles at locus i = 1,016 and locus j = 1,534, respectively. T_ij_ is the number of times that allele i and allele j occur in the same individual. A χ^2^ test was performed to determine if significant disequilibrium exists among all alleles at 1,016 and 1,534. The statistic was calculated and summed over all two-allele-interactions

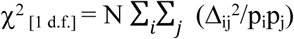

### Resistance Ratio

LC_50_ was calculated using New Orleans as a susceptible reference strain in all Resistance Ratio (RR) calculations. We calculated LC_50_ for New Orleans and for the 8 sites in generations F_3_, F_5_ and F_8_ to evaluate changes in resistance in the eight collections. Values were calculated for permethrin and deltamethrin. We used PROC CORR in SAS 9.4 to calculate Pearson’s correlation coefficient and to test for significance. This was done to examine the relationship between RR and the frequencies of I1,016 and C1,534 alleles. Analyses were performed in generations F_3_, F_6_, and F_8_ and then for all three generations combined.

## Results

### Frequency of I1,016 declines in the absence of pyrethroids

We determined the V1,016I genotype of 9,563 mosquitoes from eight locations in southern Mexico (Table 1) over eight generations (F_1_-F_8_). The V1,016I genotype counts and allele frequencies appear in Appendices 1 and 2, respectively. Figure 1A plots I1,016 frequencies in all three replicates in the eight different collections over eight generations. In general, I1,016 allele frequencies were statistically uniform among the three replicates with five exceptions as indicated with an asterisk (AcF_2_, CoF_2_, CoF_8_, DzF_2_, Mer3F_7_ - Fig. 1A).

**Fig 1.**
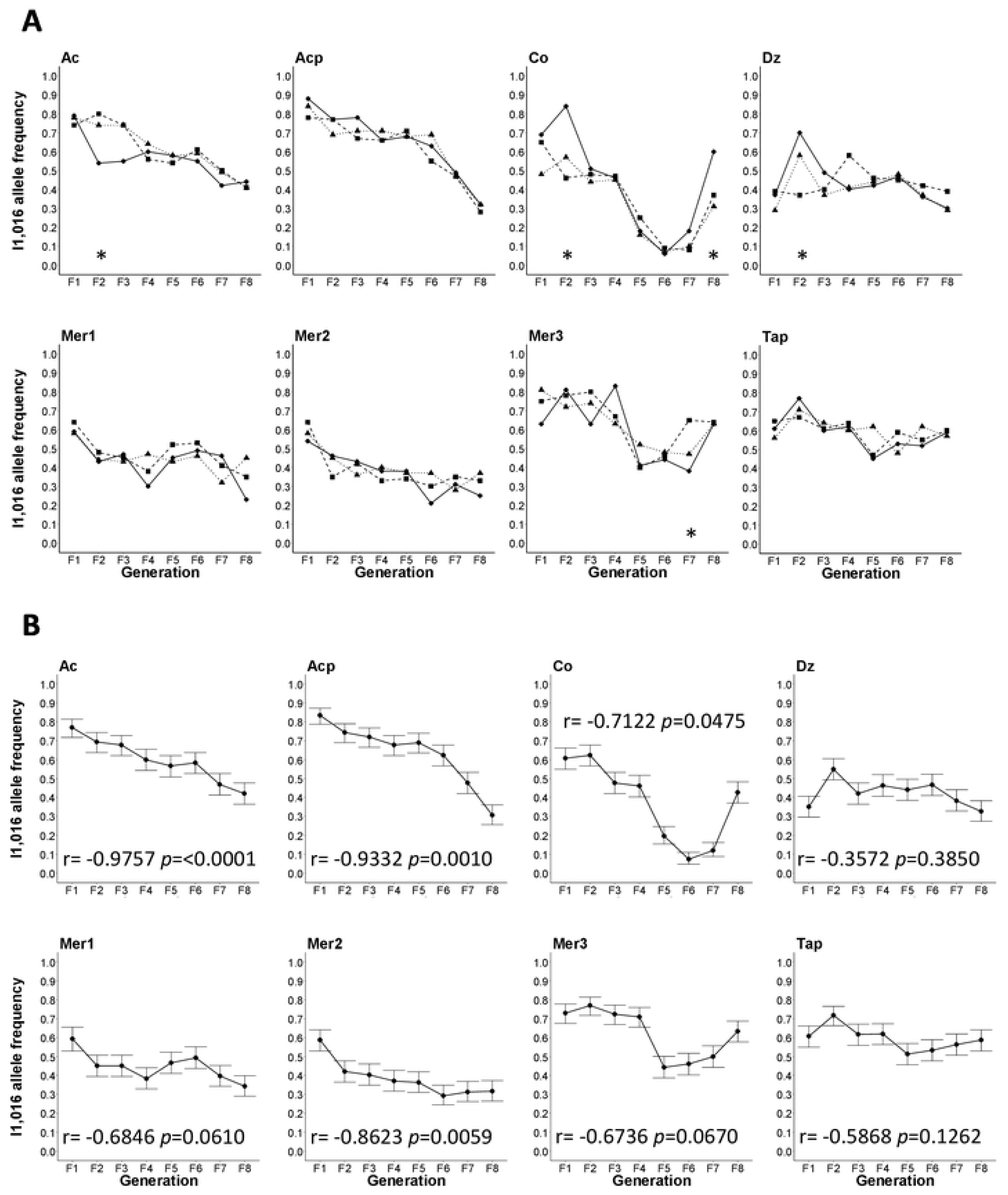
A) Plots of the frequencies of I1,016 in all three replicates in the eight different collections over eight generations. Asterisk below the three replicates at a particular generation indicate when allele frequencies were statistically different among the three replicates. B) Plots of the mean I1,016 frequencies among all three replicates and the 95% high-density intervals (HDI 95%). Pearson correlation coefficient between I1,016 frequencies and generation number and the associated significance appear at the bottom of each graph.

Figure 1B plots the mean I1,016 frequencies among all three replicates and the 95% high-density intervals (HDI 95%). Pearson correlation coefficient between I1,016 frequencies and generation number and the associated significance appear at the bottom of each graph. The correlation between I1,016 frequency and generation number was negative and significant in four out of the eight collection sites from generation F_1_ to F_8_ (Ac, Acp, Co, and Mer2) (Fig. 1B). This correlation was not significant in sites Dz, Mer1 and Tap. I1,016 frequency in Co, Mer3 and Tap declined initially and then surprisingly increased in the last 3-4 generations. In general, I1,016 declined in frequency over eight generations, however, the rate and pattern of decline varied greatly among collections.

V1,016I genotype frequencies (VV_1,016_, VI_1,016_, II_1,016_) over eight generations are plotted in Fig. 2 in each of the 8 collections. Pearson correlations coefficients and significance between genotype frequencies and generation number appear in Table 2. The correlation between VV_1,016_ frequency and generation number were all positive and four were significant.

**Table 2.** Pearson’s correlation coefficient among collection sites V1,016I genotypic frequency and generations without exposure of pyrethroids.

| Site | VV<sub>1,016</sub> |  | VI<sub>1,016</sub> |  | II<sub>1,016</sub> |  |
| --- | --- | --- | --- | --- | --- | --- |
|  | Pearson r | P value | Pearson r | P value | Pearson r | P value |
| **Ac** | 0.9301 | 0.0008 | 0.6406 | 0.0870 | -0.9418 | 0.0005 |
| **Acp** | 0.8156 | 0.0136 | 0.7455 | 0.0338 | -0.9619 | 0.0001 |
| **Co** | 0.6499 | 0.0811 | -0.4268 | 0.2916 | -0.7886 | 0.0200 |
| **Dz** | 0.2371 | 0.5718 | 0.1178 | 0.7811 | -0.4836 | 0.2247 |
| **Mer1** | 0.5472 | 0.1604 | 0.0404 | 0.9244 | -0.8005 | 0.0170 |
| **Mer2** | 0.8062 | 0.0156 | 0.3131 | 0.4502 | -0.8753 | 0.0044 |
| **Mer3** | 0.5830 | 0.1293 | 0.7033 | 0.0516 | -0.7085 | 0.0492 |
| **Tap** | 0.7559 | 0.0300 | -0.1720 | 0.6838 | -0.3615 | 0.3789 |
| **Across all** | 0.4497 | 0.0002 | 0.1462 | 0.2489 | -0.5452 | <0.0001 |

**Fig 2.**
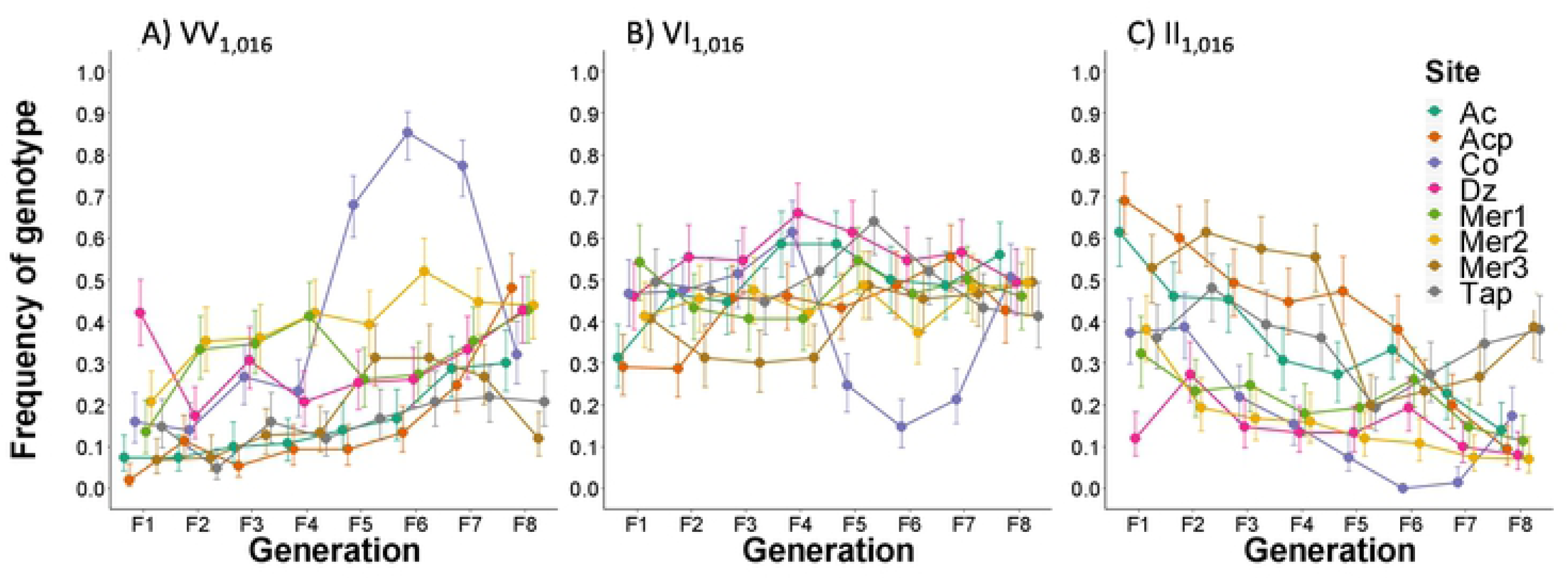
Frequencies of V1,016I genotype (VV1,016, VI1,016, II1,016) over eight generations. Six of the correlation coefficients for VI_1,016_ were positive and one was significant. All of the II_1,016_ were negative and six were significant.

Across all collections, there was a positive correlation (*r* = 0.4497, *p* = 0.0002 – Table 2) between VV_1,016_ frequencies and generation number. In contrast, the frequencies of VI_1,016_ did not change over the eight generations (*r* = 0.1462, P=0.2489 – Table 2) and were only significantly positive in Acp. II_1,016_ genotypic frequencies decreased significantly over generations (*r* = −0.5452, P < 0.0001). In general there was an overall decline in the frequency of I1,016 and an overall increase in V1,016 over eight generations.

### Frequency of C1,534 declines in the absence of pyrethroids

We genotyped F1,534C in each of the same mosquitoes for which V1,016I genotype frequencies had been determined. Appendix 1 lists the F1,534C genotypes counts, and Appendix 2 shows allele frequencies and the 95% HDI. Figure 3A shows C1,534 frequencies in the eight collection sites over eight generations. C1,534 frequencies differed among replicates in 15 of the 64 comparisons.

**Fig 3.**
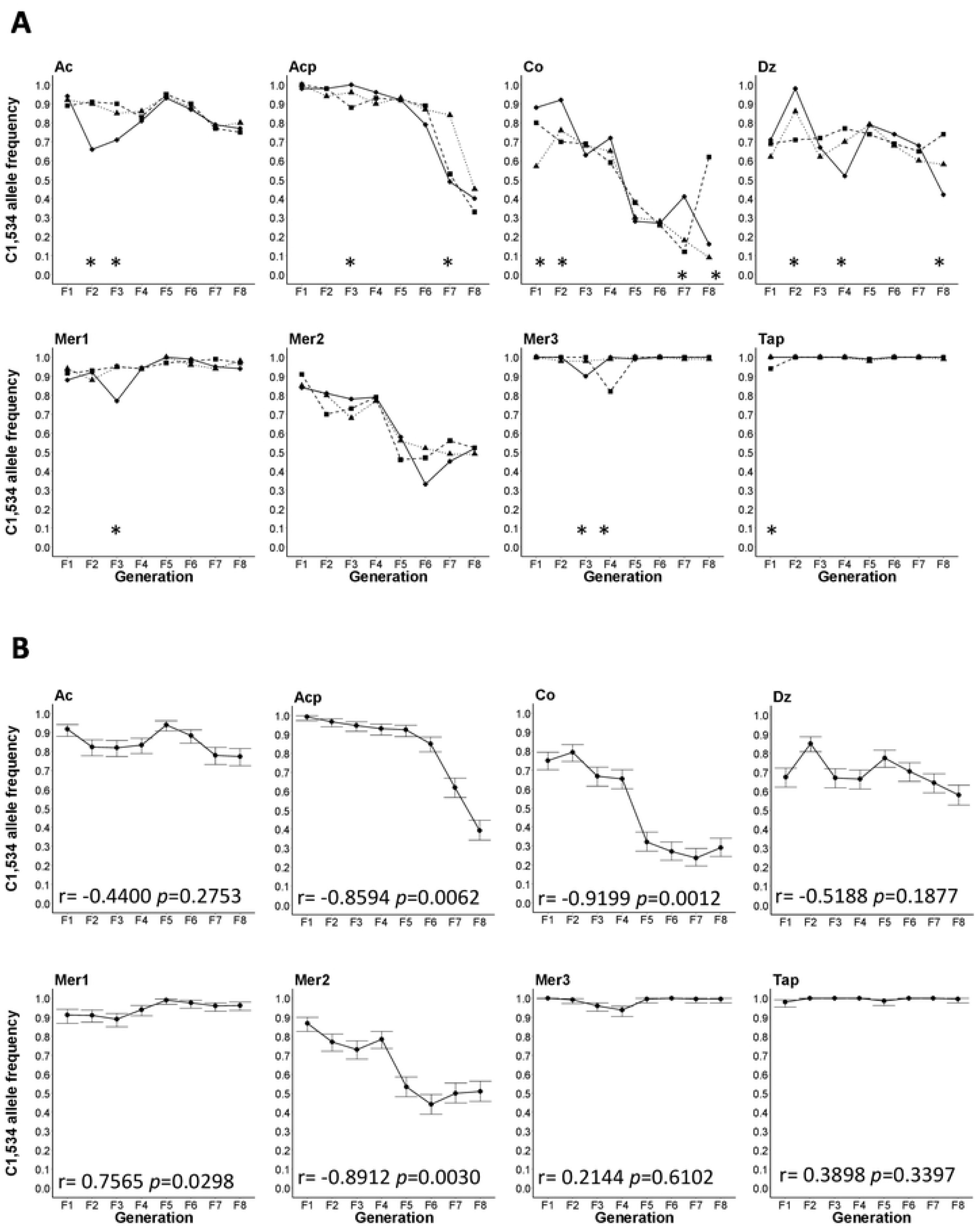
A) Plots of the frequencies of C1,534 in all three replicates in the eight different collections over eight generations. Asterisk below the three replicates at a particular generation indicate when allele frequencies were statistically different among the three replicates. B) Plots of the mean C1,534 frequencies among all three replicates and the 95% high-density intervals (HDI 95%). Pearson correlation coefficient between C1,534 frequencies and generation number and the associated significance appear at the bottom of each graph.

The correlation coefficients between C1,534 frequencies and generation number and their significance are at the base of each of the graphs in Figure 3B. The correlation coefficients between C1,534 frequency and generation number was negative and significant in four out of the eight collection sites from generation F_1_ to F_8_ (Acp, Co, Mer1, and Mer2) (Fig. 3B). The correlation was not significant in sites Ac, Dz, Mer3 and Tap. In general, C1,534 declined in frequency over eight generations, however, the rate and pattern of decline varied greatly among collections.

**Fig 4.**
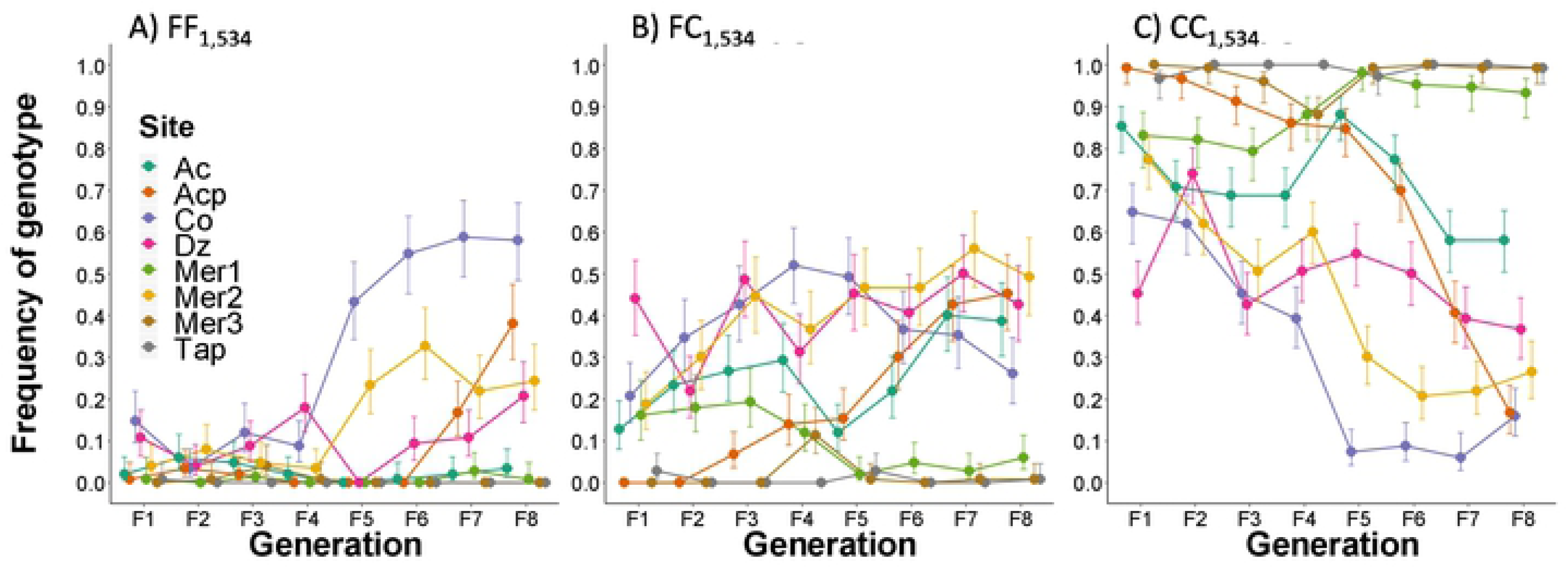
Frequencies of F1,534 genotype (FF_1,534_, FC_1,534_, CC_1,534_) over eight generations.

Correlations between genotype frequencies and generation are provided in Table 3.

**Table 3.** Pearson’s correlation coefficient among collection sites F1,534C genotypic frequency and generations without exposure of pyrethroids.

| Site | FF<sub>1,534</sub> |  | FC<sub>1,534</sub> |  | CC<sub>1,534</sub> |  |
| --- | --- | --- | --- | --- | --- | --- |
|  | Pearson r | P value | Pearson r | P value | Pearson r | P value |
| Ac | -0.363 | 0.3759 | 0.6537 | 0.0788 | -0.5460 | 0.1615 |
| Acp | 0.6952 | 0.0556 | 0.9711 | <0.0001 | -0.9110 | 0.0016 |
| Co | 0.8953 | 0.0026 | 0.0528 | 0.9012 | -0.8996 | 0.0023 |
| Dz | 0.3782 | 0.3555 | 0.3733 | 0.3624 | -0.5175 | 0.189 |
| Mer1 | 0.2729 | 0.5131 | -0.8254 | 0.0116 | 0.7953 | 0.0183 |
| Mer2 | 0.7997 | 0.0172 | 0.8748 | 0.0045 | -0.9131 | 0.0015 |
| Mer3 | -0.3423 | 0.4066 | -0.0164 | 0.9692 | 0.1308 | 0.7575 |
| Tap | -0.5774 | 0.1340 | -0.2701 | 0.5177 | 0.3309 | 0.4233 |
| Across all | 0.3399 | 0.006 | 0.2078 | 0.0994 | -0.3024 | 0.0152 |

Five correlation coefficients for FF_1,534_ were positive and two were significant, five of the correlation coefficients for FC_1,534_ were positive and three were significant and five of the CC_1,534_ were negative and four were significant. Across all collections sites, there was a positive correlation (*r* = 0.3399, *p*=0.006) between FF_1,534_ frequencies and generation number. The frequencies of FC_1,016_ did not change significantly over generations (*r* = 0.2078, P=0.0994) and as expected for a genotype that confers a lower fitness, CC_1,534_ genotypic frequencies decreased significantly over generations (*r* = −0.3024, P = 0.0152).

### Linkage disequilibrium

We performed pairwise linkage disequilibrium analyses between alleles in V1,016I and F1,534C. Table 4 lists the linkage disequilibrium coefficients R_ij_, χ^2^ and the probability value obtained between pairwise loci. Rij is distributed from −1.00 (mutations occur on opposite chromosomes - *trans*) to 0.00 (mutations occur independently), to 1.00 (both mutations *cis* on the same chromosome) and therefore provides a standardized measure of disequilibrium.

Alleles segregated in 57 out of 64 collections (Table 4). Forty seven of the 57 collections exhibited significant linkage disequilibrium, with R_ij_ values ranging between 0.15-0.85 among collections. In general and in agreement with earlier studies [13, 14] alleles in V1,016I and F1,534C were in linkage disequilibrium. Table 5 displays the correlation and the significance between haplotype frequencies and generation.

**Fig. 5.**
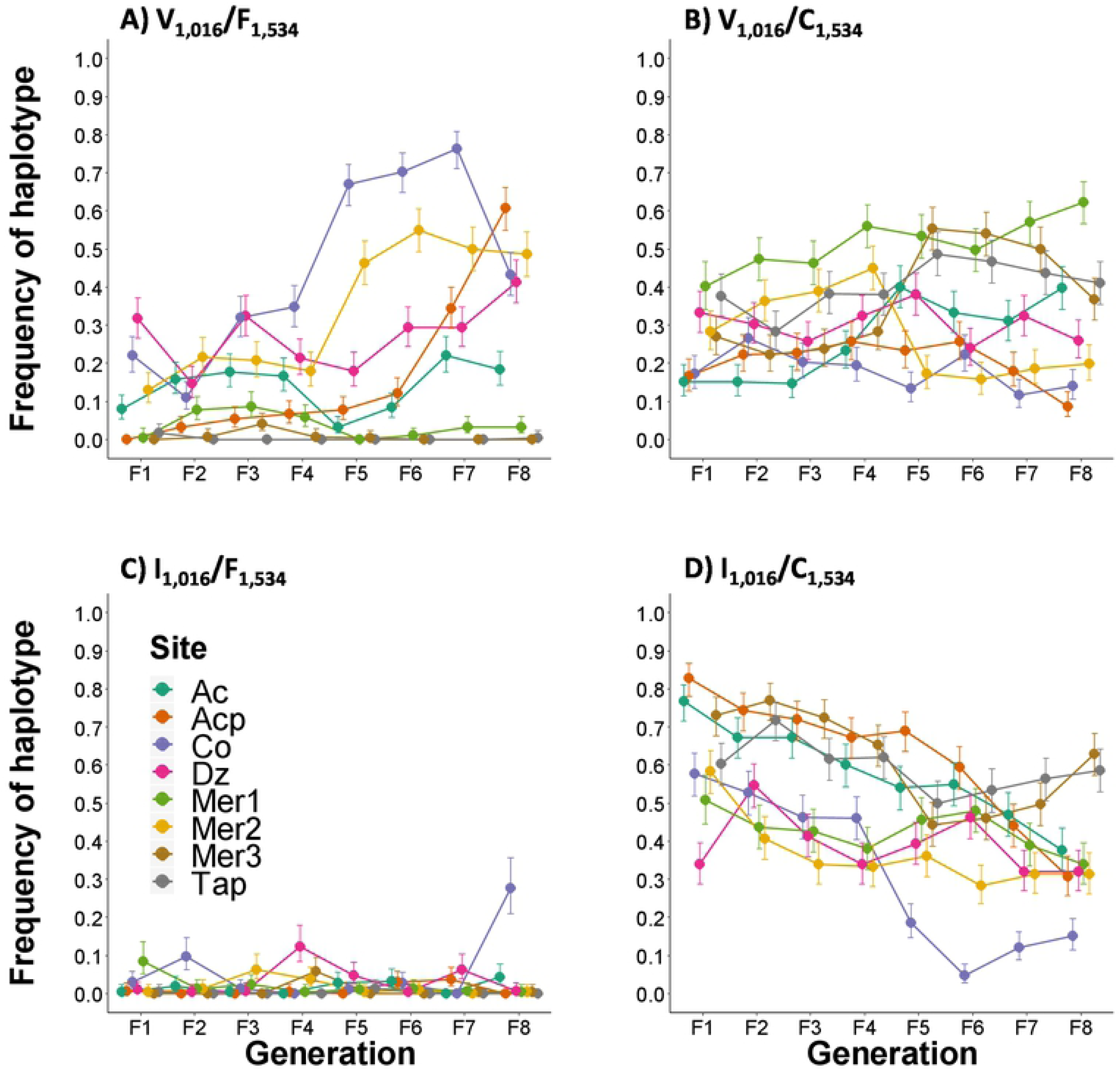
Plots the frequency of the four potential dilocus haplotypes over eight generations.

**Table 5.** Pearson’s correlation and *p* value for four haplotypes at di-locus V1016I/F1534C.

| Site | V<sub>1,016</sub>/F<sub>1,534</sub> |  | V<sub>1,016</sub>/C<sub>1,534</sub> |  | I<sub>1,016</sub>/F<sub>1,534</sub> |  | I<sub>1,016</sub>/C<sub>1,534</sub> |  |
| --- | --- | --- | --- | --- | --- | --- | --- | --- |
|  | Pearson r | P value | Pearson r | P value | Pearson r | P value | Pearson r | P value |
| **Ac** | 0.282 | 0.500 | 0.868 | 0.005 | 0.512 | 0.195 | -0.978 | <0.0001 |
| **Acp** | 0.842 | 0.009 | -0.356 | 0.386 | 0.432 | 0.285 | -0.937 | 0.001 |
| **Co** | 0.754 | 0.031 | -0.566 | 0.143 | 0.401 | 0.325 | -0.907 | 0.002 |
| **Dz** | 0.429 | 0.289 | -0.251 | 0.549 | 0.130 | 0.759 | -0.388 | 0.342 |
| **Mer1** | -0.268 | 0.521 | 0.873 | 0.005 | -0.672 | 0.068 | -0.618 | 0.103 |
| **Mer2** | 0.882 | 0.004 | -0.633 | 0.092 | -0.339 | 0.412 | -0.770 | 0.026 |
| **Mer3** | -0.340 | 0.410 | 0.677 | 0.065 | -0.031 | 0.942 | -0.688 | 0.059 |
| **Tap** | -0.480 | 0.229 | 0.624 | 0.099 | -0.051 | 0.904 | -0.560 | 0.149 |
| **Overall** | 0.323 | 0.009 | 0.140 | 0.271 | 0.091 | 0.474 | -0.516 | <0.0001 |

Table 5 displays the correlation and the significance between haplotype frequencies and generation.

The frequency of the susceptible V_1,016_/F_1,534_ haplotype increased over generations (*r* = 0.323, *p* = 0.009). The frequency of the V_1,016_/C_1,534_ haplotype remained relatively constant (*r* = 0.140, *p* = 0.271). The frequency of the resistant I_1,016_/C_1,534_ haplotype decreased over time (*r* = −0.516, *p* <0.0001) across all collection sites. The frequencies of the I_1,016_/F_1,534_ haplotypes were consistently low (*r* = 0.091, *p* = 0.474) across generations. This same trend has been noted in two previous studies [5] [15] and suggested that low fitness may occur in any mosquito in which I1,016 co-occurs with F1,534.

### Temporal analysis of di-locus genotypes

Figure 6 shows the frequency of nine di-locus genotype combinations (3 genotypes at 2 loci) and Table 6 lists the correlation between frequencies of each di-locus genotype and the generation number without pyrethroid exposure. We estimated the frequencies of the nine genotype combinations in 9,563 mosquitoes. However, the di-locus coefficient among the nine graphs was only significant for the wild type susceptible VV_1016_/FF_1534_ and the dual resistant II_1016_/CC_1534._ The correlation between generation number and VV_1016_/FF_1534_ was positive (r = 0.3376, P = 0.0064) indicating an increase while the correlation of II_1016_/CC_1534._was negative (r = −0.5465, P < 0.0001) indicating a decline. The low frequencies of VI_1,016_/FF_1,534_, II_1,016_/FF_1,534_, and II_1,016_/FC_1,534_ in Figs 6 B,C,F are consistent with the hypothesis that low fitness may occur in any mosquito in which I1,016 co-occurs with F1,534. However Figure 6E identifies an exception to this trend. A substantial proportion of the double heterozygote VI_1016_/FC_1534_ survive to generation F_8_.

**Fig 6.**
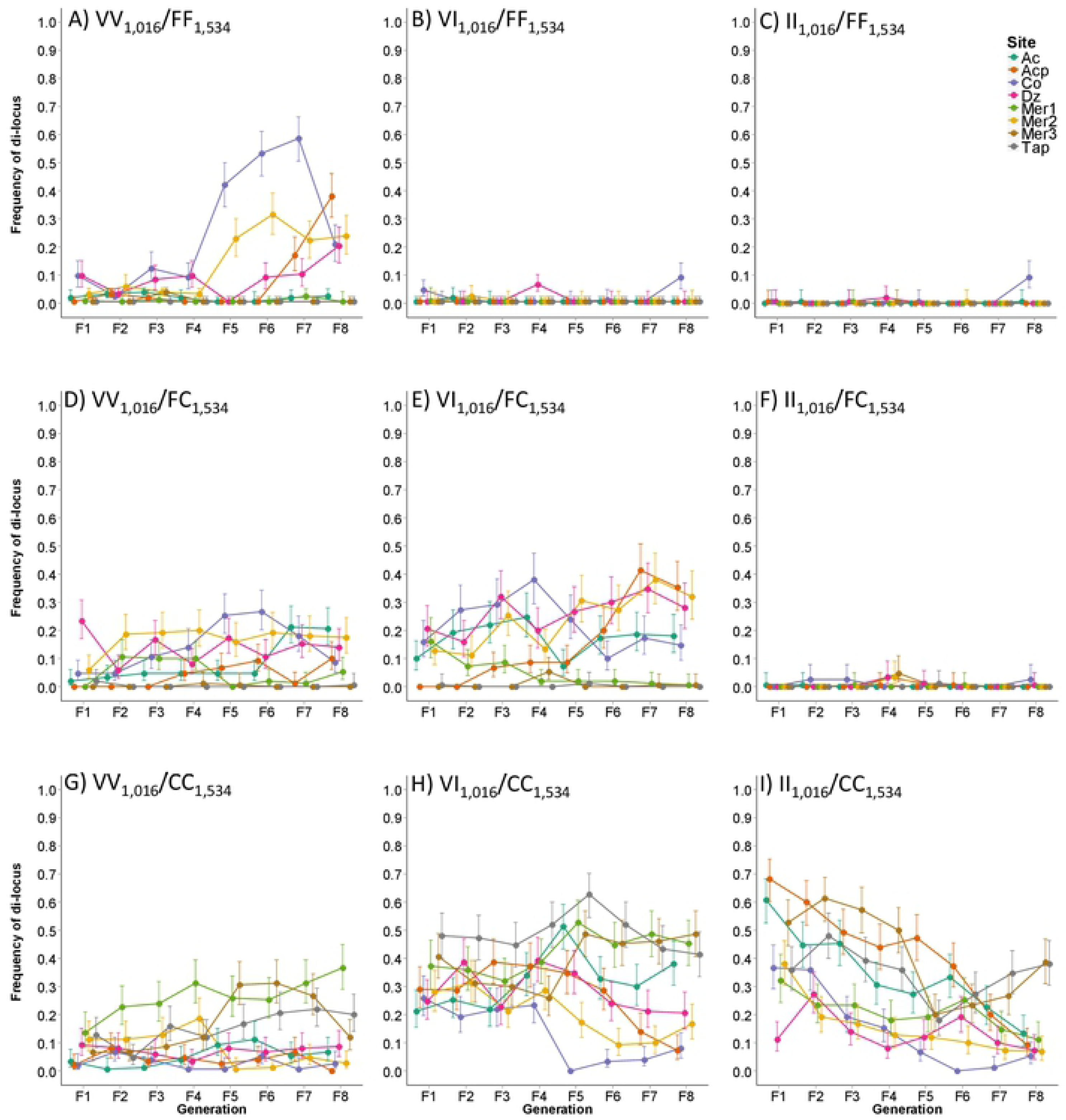
Frequency of the nine dilocus haplotypes over eight generations.

### Association between resistant allele frequencies and the Resistance Ratio

Pearson correlation coefficients and their significance were calculated between the frequencies of I1,016 or C1,534 alleles and resistance ratios for permethrin and deltamethrin as determined by bioassay for all eight collections. This was done separately for generations F_3_, F_6_, and F_8._ Table 7 indicates that all correlations were positive in F_3_ and F_8_ but none were significant. All correlations were positive in F_6_ and three were significant (**in bold**). When combining results from all three generations together, all correlations were positive and three were significant (**in bold**). Thus in general there was a weak but consistently positive correlation between the frequency of I1,016 or C1,534 alleles and resistance ratios as determined by bioassay.

**Table 7.** Pearson correlation coefficients and their significance were calculated between the frequencies of I1,016 or C1,534 alleles and resistance ratios as determined by bioassay for all eight collections. Significant values appear in bold.

| Locus | $II_{1016}$ | | | |
| --- | --- | --- | --- | --- |
| Generation | Correlation of allele frequency with Permethrin RR | Prob* | Correlation of allele frequency with Deltamethrin RR | Prob |
| $F_3$ (n=9) | 0.606 | 0.1489 | 0.625 | 0.184 |
| $F_6$ (n=9) | **0.715** | **0.0303** | **0.709** | **0.032** |
| $F_8$ (n=9) | 0.338 | 0.3732 | 0.607 | 0.083 |
| All (n=27) | **0.588** | **0.002** | **0.469** | **0.021** |

| Locus | $C_{1534}$ | | | |
| --- | --- | --- | --- | --- |
| Generation | Correlation of allele frequency with Permethrin RR | Prob | Correlation of allele frequency with Deltamethrin RR | Prob |
| $F_3$ (n=9) | 0.711 | 0.073 | 0.66 | 0.154 |
| $F_6$ (n=9) | **0.822** | **0.007** | 0.593 | 0.093 |
| F8(n=9) | 0.115 | 0.768 | 0.282 | 0.462 |
| All(n=27) | **0.707** | **<.0001** | 0.367 | 0.078 |

This was noted in an earlier study in which I1,016 showed significant protection against permethrin, and deltamethrin whereas F1534C showed protection against permethrin but not against deltamethrin [16, 17]. The expression of C1,534 in *Xenopus* oocytes and exposure to both pyrethroids demonstrated that the resistant amino acid substitution in C1,534 is sensitive to permethrin but not for deltamethrin, which is consistent with our results.

## Discussion

In this study we established eight colonies of *Ae. aegypti* from the field and maintained them in a pyrethroid free environment in the laboratory over eight generations. We demonstrated that in general the frequency of the *Ae. aegypti* pyrethroid resistance alleles I1,016 and C1,534 decline when released from pyrethroid pressure in the laboratory (Figs. 1 and 3, Tables 3 and 5). However, the pattern of decline appeared to be strain dependent with some having a steady rate of decline (Ac, Acp, Mer2 in Fig. 1, Acp and Mer2 in Fig. 3), some showing a shallow decline (e.g. Mer1, Tap in Fig 1; Acp, Mer2and Co in Fig. 3) and others displaying no net change (Fig.1 Dz; Fig 3.Ac, Mer3, and Tap).

A more surprising result was that in Co and Mer3, the frequencies of I1,016 actually increased following a precipitous drop. Likewise the frequencies of C1,534 in Mer1, Mer2, and Co, increased after a drop. This might occur if there are deleterious or lethal recessive mutations linked to the susceptible allele that became homozygous through continuous inbreeding. However this theory fails to explain why in Co I1,016 increased simultaneously in all three replicates.

We do not propose nor do we have any idea as to whether the selection pressure in indoor cage studies is the same as or is even correlated with outdoor selection pressure. This study only indicates that the loss of pyrethroid resistance is unlikely to follow a smooth linear or exponential decline for any one of a number of reasons. Nor should we expect the decline in resistance to be consistent among collections. Epistatic interactions between alleles may cause non-linear trends in allele frequencies. Much depends on the genetic background of each population; some populations could take much longer to lose resistance while others may be much faster.

There is a variety of possible causes for this variance in gene frequency trajectories among collections but it is unlikely that the heterogeneity arose from small sample sizes. We analyzed three replicates of 50 adult mosquitoes for each of the eight collection sites. Furthermore, 50 adults were used to generate the next generation of eggs for each replicate. The 95% HDI remained narrow in all graphs in Fig1B and Fig. 3B.

Initial conditions may affect the shape of the curve. For example, a curve that begins with initial frequencies close to 1 (Fig. 3A, Mer3 and Tap) would begin to decline much later than a curve that begins at 0.6 (Fig 1A, Mer1, Mer2). Metabolic resistance may account for much of this heterogeneity. A QTL mapping study [9] reported that 58.6% of the variation in knockdown could be accounted for by I1,016 but that a number of different QTL located throughout the genome accounted for the remainder of the variation. Saavedra-Rodriguez et al used the ’Aedes Detox’ microarray [18] and showed an inverse relationship between I1,016 frequencies and the numbers of differentially transcribed metabolic genes[19].

Table 5 displays the correlation and the significance between haplotype frequencies and generation. The low frequencies of VI_1,016_/FF_1,534_, II_1,016_/FF_1,534_, and II_1,016_/FC_1,534_ noted in this (Fig 6 B,C,F) and two previous studies [5], [15] suggests that low fitness may occur in any mosquito in which I1,016 co-occurs with F1,534. This and the fact that C1,534 historically appears before I1,016 argues that evolution of the mutations were sequential. If I1,016 had appeared first it would have co-occured with F1,534 and would have been eliminated.

Deltamethrin RR appears to be correlated with I1,016 allele frequency but not with C1,534 allele frequencies, while permethrin RR correlates with both allele frequencies (Table 7). We speculated that there are other resistant mechanisms that could drive resistance to deltamethrin, such as detoxifying enzymes or other mutations in *VGSC* that have not been identified. However, permethrin resistance appears to be driven heavily by both resistance alleles.

The I1,016, amino acid substitution has been found in many resistant populations in the Americas [20–25] and it has been shown that it is in linkage disequilibrium with C1,534 [12]. Recent work has shown that I1,016 is also in very tight disequilibrium with V410L [16].

Interestingly, the I1,016 mutation was functionally expressed in *Xenopus* oocytes and did not show an alteration to the sodium channel sensitivity to both pyrethroids [16]. Our data indicate a positive correlation of I1,016 allele frequency with the RR of both pyrethroids. It is clear that more understanding is needed as to the role that I1,016 plays in resistance of *Ae. aegypti.* This includes the possibility that V410L [16] may be the residue to which the pyrethroid actually binds.

Since many field populations are resistant to pyrethroid and few insecticides are available in the market, negative fitness is beneficial for vector control, providing an opportunity where alternative insecticide could be used while losing pyrethroid resistance. Pyrethroids could then be “saved” to control susceptible populations during a disease outbreak. However it is important to remember that even if dominant wild type alleles (V1,016 and F1,534) increase in a population, recessive resistance alleles will be hidden and maintained. There is no way that the initial susceptibility of populations (before the introduction of a pyrethroid) will ever be recovered. Recent work in Sao Paulo State in Brazil showed that resistance alleles persist in natural populations for at least 11 years [1].

## Acknowledegments

This study was funded by the National Institutes of Health/National Institute of Allergy and Infectious Diseases International Collaborations in Infectious Disease Research Program (U01-AI-088647) and “Insecticide Resistance Management to Preserve Pyrethroid Resistance in Aedes aegypti” (1R01AI121211-01A1). Farah Vera-Maloof and Patricia Penilla were supported by the Fogarty Training Grant “Training in Dengue Prevention and Control” (2D43TW001130-08).

